# *Salmonella* effector SseL induces PD-L1 up-regulation and T cell inactivation via β-catenin signalling axis

**DOI:** 10.1101/2024.10.29.620790

**Authors:** Umesh Chopra, Maria Kondooparambil Sabu, Raju S Rajmani, Ayushi Devendrasingh Chaudhary, Shashi Kumar Gupta, Dipshikha Chakravortty

## Abstract

The upregulation of PD-L1 by various pathogens is a recognized strategy to evade the adaptive immune response. *Salmonella* infection also upregulates PD-L1 levels causing culling of the activated T-cell; however, the underlying mechanism behind this upregulation is not known. Our findings indicate that the upregulation of PD-L1 is through *Salmonella* pathogenicity island 2 (SPI-2) encoded effectors since PFA-fixed STM WT and STM*ΔssaV* (which is unable to secrete effector proteins) did not alter PD-L1 levels. We have further investigated the role of the SPI-2 effector SseL (a deubiquitinase known to affect the NF-ĸB pathway) in PD-L1 upregulation. Our study identifies SPI-2 effector SseL to be crucial for upregulating PD-L1 *in vitro* as well as *in vivo* murine models. The increase in PD-L1 levels induced by STM WT facilitates colonization in secondary infection sites in C57BL/6 mice, including the liver and spleen, while the STM*ΔsseL* strain exhibits significant colonization defects. Notably, despite the reduced colonization capacity of STM*ΔsseL*, infected mice exhibit earlier mortality associated with heightened inflammation. We further elucidated the molecular mechanism behind PD-L1 upregulation and observed that bacterial effector SseL helps in the stabilization of β-catenin inside the cell. β-catenin thus translocates into the nucleus and directly regulates the transcriptional levels of PD-L1, which is abrogated upon using β-catenin/TCF inhibitor FH535. Collectively, our study elucidates the mechanism by which *Salmonella* mediates immune suppression through PD-L1 upregulation.

Abstract figure: Schematic representation of SseL mediated PDL1 upregulation and further affecting the T cell proliferation

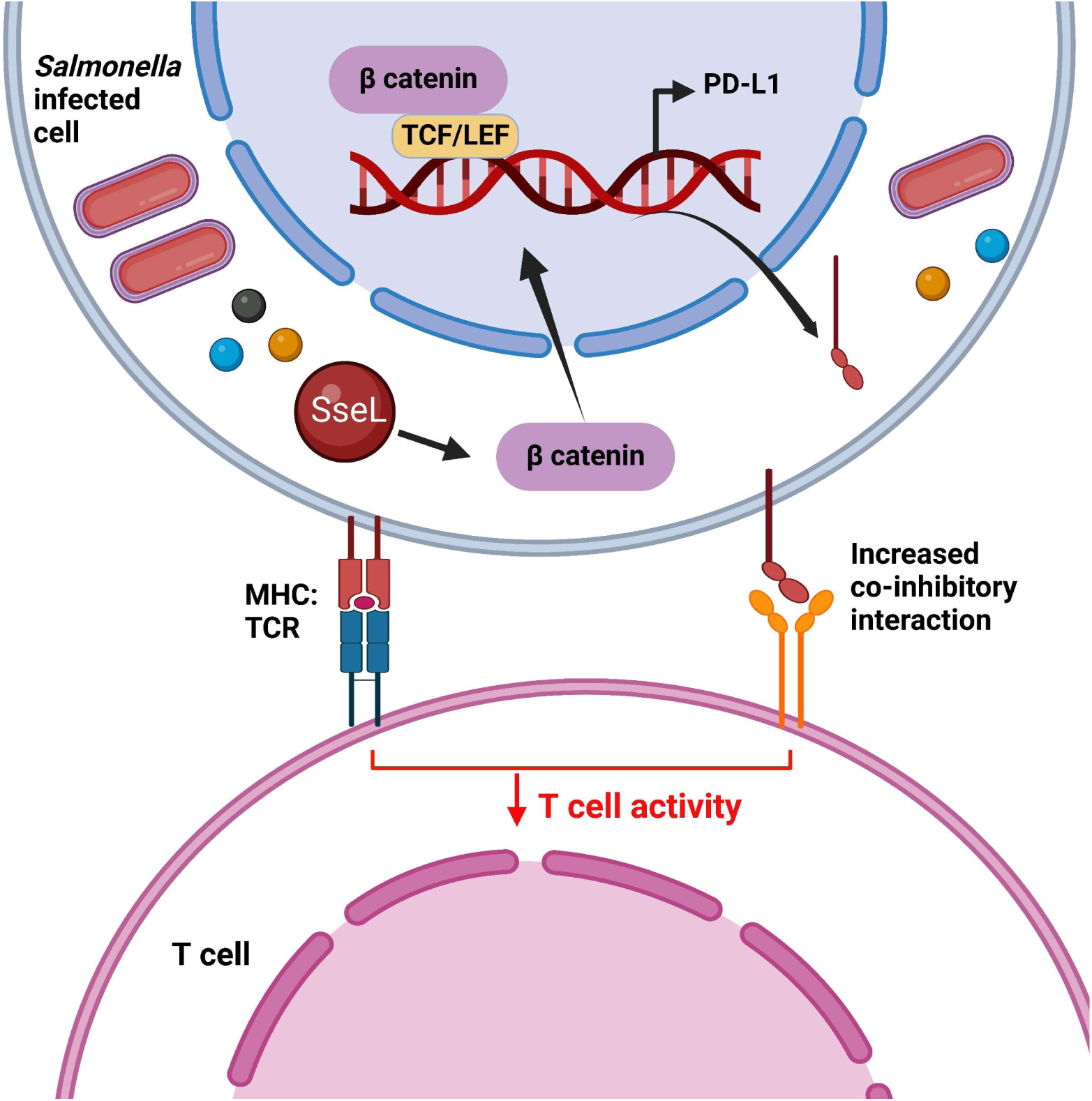

## Introduction

*Salmonella* is an enteric pathogen that proliferates and survives within host cells. It infects a wide range of hosts and manifests disease with varied symptoms, ranging from gastroenteritis to severe systemic infections[1]. As per WHO reports, *Salmonella* is one of the four leading causes of diarrhoeal diseases in humans, with typhoid fever, caused by *Salmonella* Typhi, accounting for nearly 9 million infections and 110,000 deaths in 2019[2], [3], [4]. *Salmonella* enters the human body through the ingestion of contaminated food and water and reaches the intestinal lumen. It invades the intestinal epithelium via transcytosis through specialized microfold cells (M cells) overlying the Peyer’s Patches and reaches the underlying lamina propria [1]. The pathogen also invades non-phagocytic intestinal epithelial cells through the action of effector proteins injected by the Type 3 secretion system (T3SS)-1 encoded by *Salmonella* pathogenicity island-1 (SPI-1)[5], [6], [7]. Effectors injected by T3SS-2 encoded by SPI-2 aid the pathogen in intracellular survival and proliferation[8].

Macrophages, dendritic cells, and neutrophils in the lamina propria mount the early innate immune response against *Salmonella* through phagocytosing free bacteria and releasing pro-inflammatory cytokines such as TNF-α[9]. They also present *Salmonella* antigens on MHC molecules for recognition by T cell receptors (TCRs) on T cells[10]. *Salmonella* clearance from host tissues depends majorly on the functionality of CD4^+^ T cells with a contribution of CD8^+^ T cells in the later stages of infection[11], [12]. *Salmonella* secretes a plethora of effectors aimed at modulating the functions of T cells to prolong colonization and survival in the host. *Salmonella* effector SifA interferes with surface expression of MHC II on APCs[13], while SteD mediates MARCH8-dependant ubiquitination of MHC II molecules, suppressing T cell activation[14], [15].

PD-L1(Programmed death ligand-1) is a hallmark immuno-regulatory ligand belonging to the B7 family, present on both hematopoietic and non-hematopoietic cells, expressed constitutively or upregulated during inflammation[16]. PD-L1 binds to its receptor, PD-1, on activated T cells, leading to inactivation of T cells, presented by decreased inflammatory cytokine production, T-cell proliferation, and survival[17]. The PD-1: PD-L1 co-inhibitory interaction is an essential component of peripheral tolerance and plays a vital role in controlling inflammation[18], [19]. *Salmonella* is known to facilitate up-regulation of PD-L1 in B cells [20] and intestinal epithelial cells [21] leading to anergy of activated CD4^+^ T cells and CD8^+^ T cells [22], [23] resulting in reduced pathogen clearance. This is a successful immune evasion strategy employed by several other pathogens including parasites (*Schistosoma mansoni*), bacteria (*Mycobacterium tuberculosis, Helicobacter pylori*), and viruses (Hepatitis B virus, Lymphocytic choriomeningitis virus)[24], [25], [26], [27], [28], [29], [30], [31].

In the case of *H. pylori* infection, bacterial effector CagA-dependent PD-L1 expression is known, while the surface proteins of *Porphyromonas gingivalis* and *Streptococcus pneumoniae* upregulates PD-L1 expression [24], [32][33]. During *Salmonella* infection, a dominant role of *Salmonella* pathogenicity island-2 [23] and certain SPI-2 effector proteins are known [21] in the up-regulation of PD-L1 in intestinal epithelial cells; however, the molecular mechanism behind this phenomenon remains elusive.

Several cancers also upregulate PD-L1 expression to evade the immune response. In case of glioblastoma, the aberrant signalling of the Wnt/β catenin pathway has been implicated in the up-regulation of PD-L1 [34]. In the present study, we demonstrated that in the case of *Salmonella* infection, the effector protein SseL plays a crucial role in PD-L1 up-regulation *in vivo* as well as *in vitro*. SseL modulates β-catenin in the cytosol, and β-catenin directly regulates the level of PD-L1 expression. Compared to STM WT infected, mice infected with STM*ΔsseL* showed reduced organ burden, but a severe inflammatory response characterized by histopathology analysis and quantification of pro-inflammatory cytokines. These results suggest a role for PD-L1 in controlling inflammation in due course of *Salmonella* infection and colonization of the pathogen. SseL proves to be a potential therapeutic target due to its ability to modulate immune responses in favour of the *Salmonella* infection.

### Experimental procedures

#### Cell Culture and Maintenance

RAW 264.7 murine macrophages, Caco2, and bone marrow-derived macrophages (BMDM) were cultured in DMEM (Lonza) containing 10% FBS (Gibco) in a humified incubator maintaining a temperature of 37°C and 5% CO_2_. Cells were seeded in 6 wells, 12 wells, or 24 well plates at a confluency of 60-70% prior to infection. Cells were maintained in the presence of 1% penstrep. Cells were given a PBS wash and supplemented with fresh media without penstrep at least 6 hours prior to infection. BMDM were isolated from 6–8-week-old Balb/C mice by using the published protocol [35].

#### Bacterial strains and culture conditions

*Salmonella enterica* subspecies *enterica* serovar Typhimurium (STM WT) wild-type strain ATCC 14028s was used in all experiments. The bacterial strain was cultured in Luria broth (LB-Himedia) with constant shaking (170 rpm) at 37°C orbital-shaker. Antibiotics like kanamycin (50µg/ml), and ampicillin (50µg/ml) were used wherever required. A one-step gene inactivation method was used for the generation of knockout strains [36]. Wild type as well mutant strains were transformed with pFPV-m-cherry plasmid for immunofluorescence assays.

**Table 1:** List of bacterial strains used in this study.

| Strain | Characteristic<br>(Antibiotic<br>resistance) | Source |
| --- | --- | --- |
| STM WT 14028s | - | Kind gift by Prof. M. Hensel |
| STMΔ*ssaV* | Kan<sup>R</sup> | Laboratory stock |
| STMΔ*sseL* | Kan<sup>R</sup> | This study |
| STMΔ*sseL::sseL* | Kan<sup>R</sup> +Amp<sup>R</sup> | This study |
| STM WT expressing mCherry | Amp<sup>R</sup> | Laboratory stock |
| STMΔ*sseL* expressing mCherry | Kan<sup>R</sup> +Amp<sup>R</sup> | This study |

#### Generation of Complement strain

Low copy number plasmid pQE-60 was used to prepare the STM*ΔsseL:sseL* complement strain. Briefly, PCR amplification (using Thermo Q5 DNA polymerase) of the *sseL* gene was performed using Wild Type STM colony with respective PCR primers. Forward cloning primer was chosen from a gene start site. The amplified PCR product was purified using a Gel extraction kit (Qiagen). The PCR product and Plasmid were subjected to restriction digestion by using Xho1 (NEB) and Hind III (NEB) at 37°C for 2 hours. Double-digested PCR product and plasmid were further subjected to Ligation using T4 DNA ligase (NEB) at 16°C for 12-14 hours. The ligated product was further electroporated in STMΔ*sseL* strain to prepare the complemented strain i.e. STM*ΔsseL*:pQE60-*sseL* (hence referred to as STM*ΔsseL:sseL*). The expression level of complement was confirmed with western blotting and confirmed with DNA sequencing.

#### Protocol for infection (Gentamicin Protection Assay)

The cells were seeded at the required confluency and infected with *Salmonella* strains with actively growing log phase culture (for epithelial cells) and stationary phase culture (for RAW 264.7, BMDM macrophages) at Multiplicity of infection (MOI) 10, for western blot and qRT-PCR analysis, and MOI 50 for Caco2 infection for western blot, immunofluorescence and qRT-PCR. For confocal microscopy studies, cells were seeded on sterile glass coverslips at least 24 hours before infection. Upon infecting the cells at the required MOI, the plate was centrifuged at 600 rpm for 5 min to facilitate the adhesion, and then the plate was incubated for 25 min at 37°C and 5% CO_2_. Post incubation, bacteria-containing media was removed, and cells were given 2 PBS washes to remove any extracellular bacteria. Fresh media containing 100µg/ml of gentamicin was added into wells and incubated at 37°C for one hour. Following this, cells were again given 2 PBS washes, and fresh media containing 25µg/ml of gentamicin was added, and this concentration was maintained till the end point of the experiment. Cells were harvested 24 hours post-infection.

#### Confocal Microscopy

24-hour post-infection, cells were washed with PBS twice and fixed with 3.5% PFA at room temperature for 15 min. After fixation, coverslips were washed with PBS, and cells were stained with primary antibody diluted in 2% BSA in PBS carrying 0.01% of saponin (for membrane permeabilization) for 1 hour at room temperature or 4°C overnight in a cold room as required. Post incubation, coverslips were washed again twice with PBS and incubated with specific secondary fluorophore-labelled antibodies and incubated for one hour at room temperature. DAPI was used to stain the nucleus. Coverslips were then mounted on clean glass coverslips using mounting media. Once the mounting media was dried, coverslips were stabilized by sealing the periphery with transparent nail paint on the sides. Images were acquired on a confocal scanning laser microscope (Zeiss 880 multiphoton microscope). Calculations of CTCF were performed using ImageJ for intensity measurement. Antibodies used in this study are PD-L1 (CST 13684S), β-catenin (CST 9562S), p-β-catenin (CST 9561T), and LAMP1 (DSHB 1D4B).

#### RNA isolation and Real-Time PCR (qRT-PCR)

RNA isolation was performed using TRIzol (Takara) following the manufacturer’s protocol. Quantification of RNA was done using nano-drop (Thermo-Fisher Scientific) and RNA samples were run on 1% agarose gel to check the quality of RNA. 2 µg of RNA was subjected to DNase I (Thermo Fischer Scientific) treatment at 37°C for 1 hr followed by addition of 0.5M EDTA (final concentration 5mM) and heat inactivation at 65°C for 10 mins to inactivate the enzyme. Further cDNA was prepared by using a cDNA synthesis kit (Takara) as per the manufacturer’s instructions. The expression profile of all genes of interest was evaluated using RT primers by using SYBR green mix (Takara) in a Biorad Real-Time PCR machine. The expression value of the target gene was normalized to housekeeping internal control β-actin and further compared with uninfected cells.

#### *In vivo* animal experiment

6-8 weeks old C57BL/6 male mice were used in this study. 5 days post-infection, mice were euthanized, and organs such as liver, spleen, MLN, intestine, and blood were harvested to study the colonization in organs. Organs were homogenized and further plated on *Salmonella*-*Shigella* agar to enumerate CFU. Obtained CFU values were normalized with the gram weight of the organ and then converted into a log scale. Blood was collected through heart puncture and used for ELISA. For the mice survival assay, 10^8^ CFU was administered to each mouse via oral gavage. The mice were monitored daily until all individuals in the cohort had succumbed. The animal experiments were carried out in accordance with the approved guidelines of the institutional animal ethics committee at the Indian Institute of Science, Bangalore, India (Registration No: 48/1999/CPCSEA). All procedures involving the use of animals were performed according to the Institutional Animal Ethics Committee (IAEC)-approved protocol.

#### ELISA

Quantitative estimation of pro-inflammatory cytokines was performed through ELISA as per the manufacturer’s protocol (KRISHGEN BioSystems). Briefly, 96 well plates were incubated with mice serum for 1-2hrs at room temperature. Following this, plates were washed 3 times with wash buffer, and 100μl of detection antibody was added and incubated for 1 hour at room temperature. Post incubation, the plate was washed three times with wash buffer and further incubated with Streptavidin:HRP solution and incubated for 15 min. Post incubation, the plate was washed again, and TMB substrate was added for 30 min at room temperature. Later, stop solution was added, and absorbance was captured at 450nm.

#### Chromatin Immunoprecipitation (ChIP) Assay

Post-infection, to crosslink the DNA-Protein interactions of the cells, Caco2 cells were fixed with 3.7% formaldehyde (15min, RT) followed by the addition of 125 mM glycine to quench the reaction. Lysis of the nuclei was carried out in 0.1% SDS lysis buffer [50 mM Tris-HCl (pH8.0), 200 mM NaCl, 10 mM HEPES (pH 6.5), 0.1% SDS, 10mM EDTA, 0.5 mM EGTA, 1 mM PMSF, 1 μg/ml each of aprotinin, leupeptin, pepstatin, 1 mM Na3VO4 and 1 mM NaF] followed by shearing of chromatin using Bioruptor Plus (Diagenode) at high power for 70 rounds of 30 sec pulse ON/45 sec OFF. Chromatin extracts with DNA fragments of an average size of 500 bp were immunoprecipitated using a specified antibody or anti-rabbit IgG complexed with Protein A agarose beads (Bangalore Genei). Further, Immunoprecipitated complexes were sequentially washed thrice in each buffer [Wash Buffer A: 50 mM Tris-HCl (pH8.0), 500 mM NaCl, 1 mM EDTA, 1% Triton X-100, 0.1% Sodium deoxycholate, 0.1% SDS and protease/phosphatase inhibitors; Wash Buffer B: 50 mM Tris-HCl (pH 8.0), 1 mM EDTA, 250mM LiCl, 0.5% NP-40, 0.5% Sodium deoxycholate and protease/phosphatase inhibitors; TE: 10 mM Tris-HCl (pH 8.0), 1 mM EDTA] and eluted in elution buffer [1% SDS, 0.1 M NaHCO3]. RNA and Protein were degraded from the eluted samples with RNase A and Proteinase K, and DNA was precipitated using the phenol-chloroform-ethanol extraction method. Analysis of the purified DNA was carried out by quantitative real time PCR. All values in the test samples were normalized to amplification of the specific gene in Input and IgG pull down and represented as fold change in modification or enrichment.

#### *In vivo* knockdown

For *in vivo* knockdown, adeno-associated virus serotype 2 (AAV2) was utilized. AAV2 was produced in HEK293T cells. Initially, HEK293T cells were transfected with an AAV plasmid containing shRNAs targeting PD-L1 (sequence: CUUCUGAGCAUGAACUAAU) and a scramble control under the U6 promoter, along with a helper plasmid using PEI Max 40000 (Polysciences USA). The following day, the transfection medium was replaced with a fresh culture medium. After 48 hours, the medium was collected, and fresh culture medium was added for another 48 hours. The culture medium was collected again, and the cells were harvested via trypsinization. The collected medium, containing secreted AAV viral particles, was incubated with 40% PEG 8000 overnight. The precipitated viral particles were collected by centrifugation and re-suspended in PBS. The cells containing AAV viral particles were lysed with citrate buffer and treated similarly with 40% PEG 8000 for viral precipitation. The precipitated particles were cleaned with chloroform and loaded onto an iodixanol gradient for further purification using ultracentrifugation. The cleaned viral particles were collected from the 40% gradient and transferred to Amicon 100K cut-off columns. The purified AAV particles were then titrated using real-time PCR. For AAV2 infection, mice were retro-orbitally injected with 200 μl containing approximately 2*10^12^ viral particles with the respective scrambled or shRNA constructs. On the fourteenth-day post-injection, the mice were orally infected with 10^7^ CFU of *S*. Typhimurium. Five days after infection, the mice were euthanized, and organs were collected. The knockdown was confirmed by performing western blotting on the harvested liver tissue (Fig: S2A, B).

#### Luciferase assay

For the luciferase assay, Caco2 cells were seeded in 12-well plates. Eighteen hours after seeding, the cells were transfected with luciferase construct (M50 Super 8X TOPFlash, addgene plasmid#12456, kind gift from Prof. Kumaravel Somasundaram, IISc) using lipofectamine 3000 (transfection reagent). After 48 hours, the cells were infected with STM WT and STM*ΔsseL*, while uninfected cells served as control for 24 hours. Following the infection period, cells were harvested and lysed in reporter lysis buffer (Promega). An equal amount of protein was used for the luciferase assay.

### Statistical analysis

Statistical analysis was performed using GraphPad Prism software. For analysis, a one-way ANOVA test was performed to obtain the p values. (****p<0.0001, ***p < 0.001, ** p<0.01, *p<0.05). The results are indicative of mean± SD or mean± SEM. For animal experiments, the Mann-Whitney test was performed. Group size, experimental number, no of cells in each set, and P values of each experiment are described in figure legends.

**Table 2:**
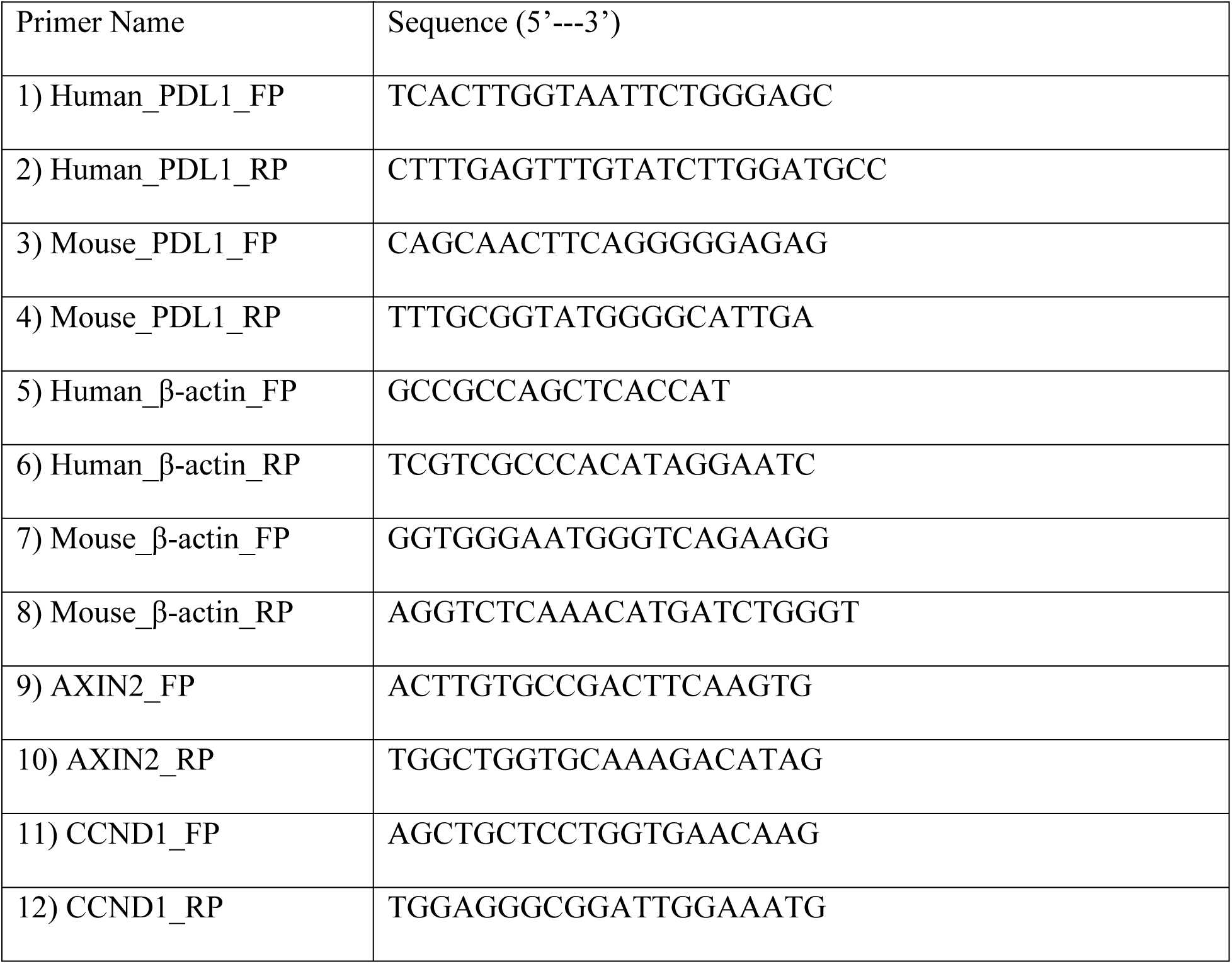
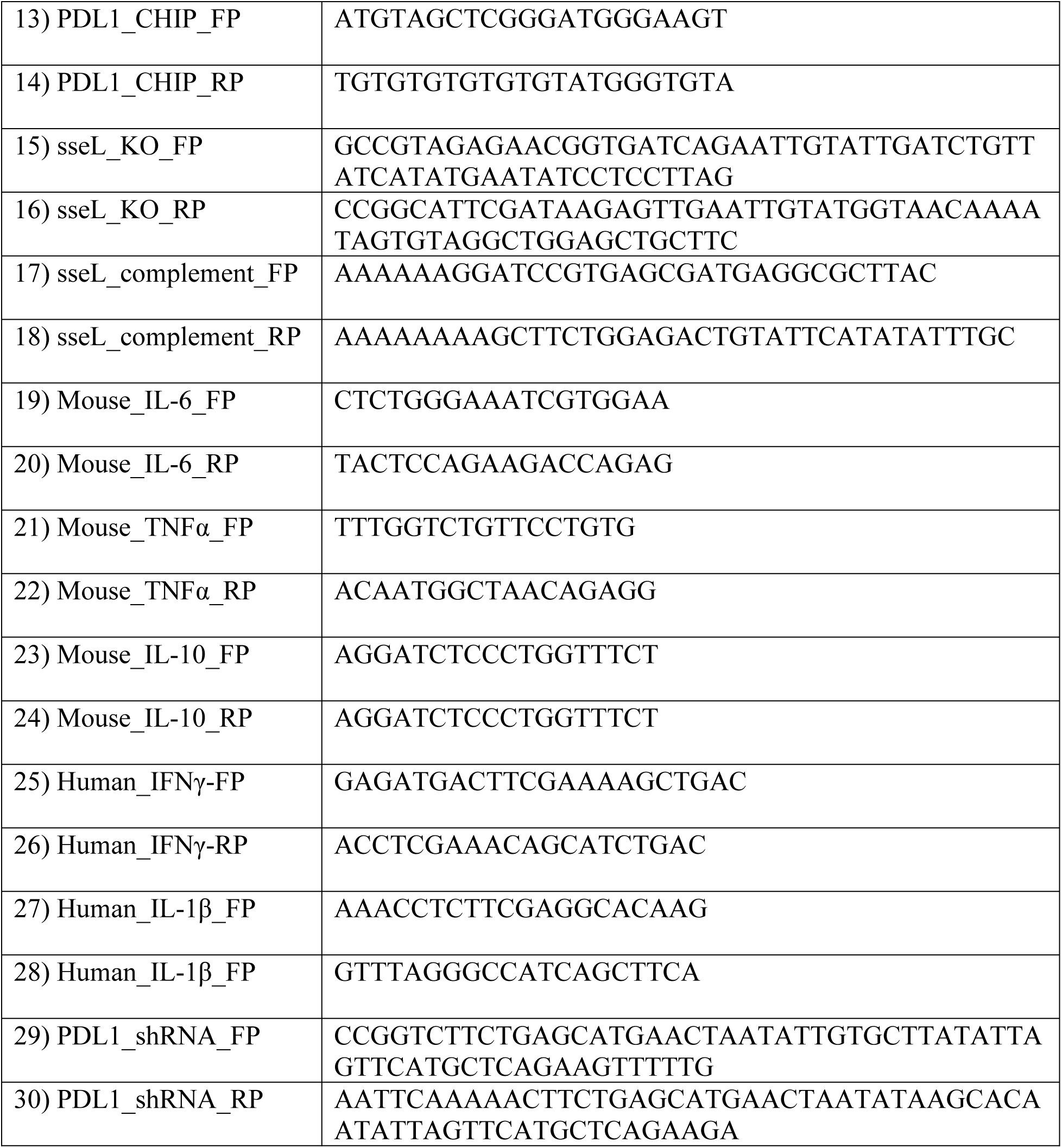
Primer sequence used in study.

## Results

### *Salmonella* upregulates the level of PD-L1 *in vitro* as well as *in vivo* using *Salmonella* pathogenicity island 2 (SPI-2)

Several viral and bacterial species are known to upregulate the level of PD-L1 either through the action of effector proteins or by utilizing surface PAMPs for their own survival. Recent studies suggest that upon *Salmonella* infection, there is an increase in PD-L1 level. To investigate if, in the case of *Salmonella* infection, PAMPs play any role in PD-L1 upregulation, we have infected the RAW 264.7, BMDM, and Caco2 cell line with *Salmonella* WT 14028s (mentioned as STM WT) or STM WT 14028s strain fixed with PFA (which maintains intact PAMPs) and 24 hours post-infection, quantified the transcript level of PD-L1. Our data suggests that compared to uninfected (UI) and STM WT (PFA fixed) infected cells, there is a significant increase in the level of PD-L1 upon infection with STM WT in RAW264.7, BMDM, and Caco2 cells respectively, suggesting that PAMPs are not crucial for the upregulation of PD-L1 while STM WT using effector proteins can upregulate the PD-L1 level (Fig-1 A, B and C). To further elucidate the importance of *Salmonella* effectors in regulating the PD-L1 levels, 6–8-week-old C57BL/6 mice were infected with STM WT or STM*ΔssaV* (mutant strain which is unable to secrete SPI2 encoded effector from SCV to host cytosol) via oral gavage while PBS treated cohort was kept as a control. 5^th^ day post-infection, mice were euthanized, and organs like the liver, spleen, and intestine were harvested for analysis. Quantification of data shows a marked increase in PD-L1 at transcript level upon STM WT infection compared to PBS-treated and STM*ΔssaV* infected in organs like the liver, spleen, and intestine (Fig-1 D, E, and F). Together, this data suggests that upon *Salmonella* infection, there is an increase in the level of PD-L1 *in vitro* as well as *in vivo*, and this upregulation is dependent on *Salmonella* SPI2 encoded effector proteins as demonstrated by the inability of STM WT-PFA-fixed and STM*ΔssaV* strains to enhance PD-L1 levels.

**Fig 1:**
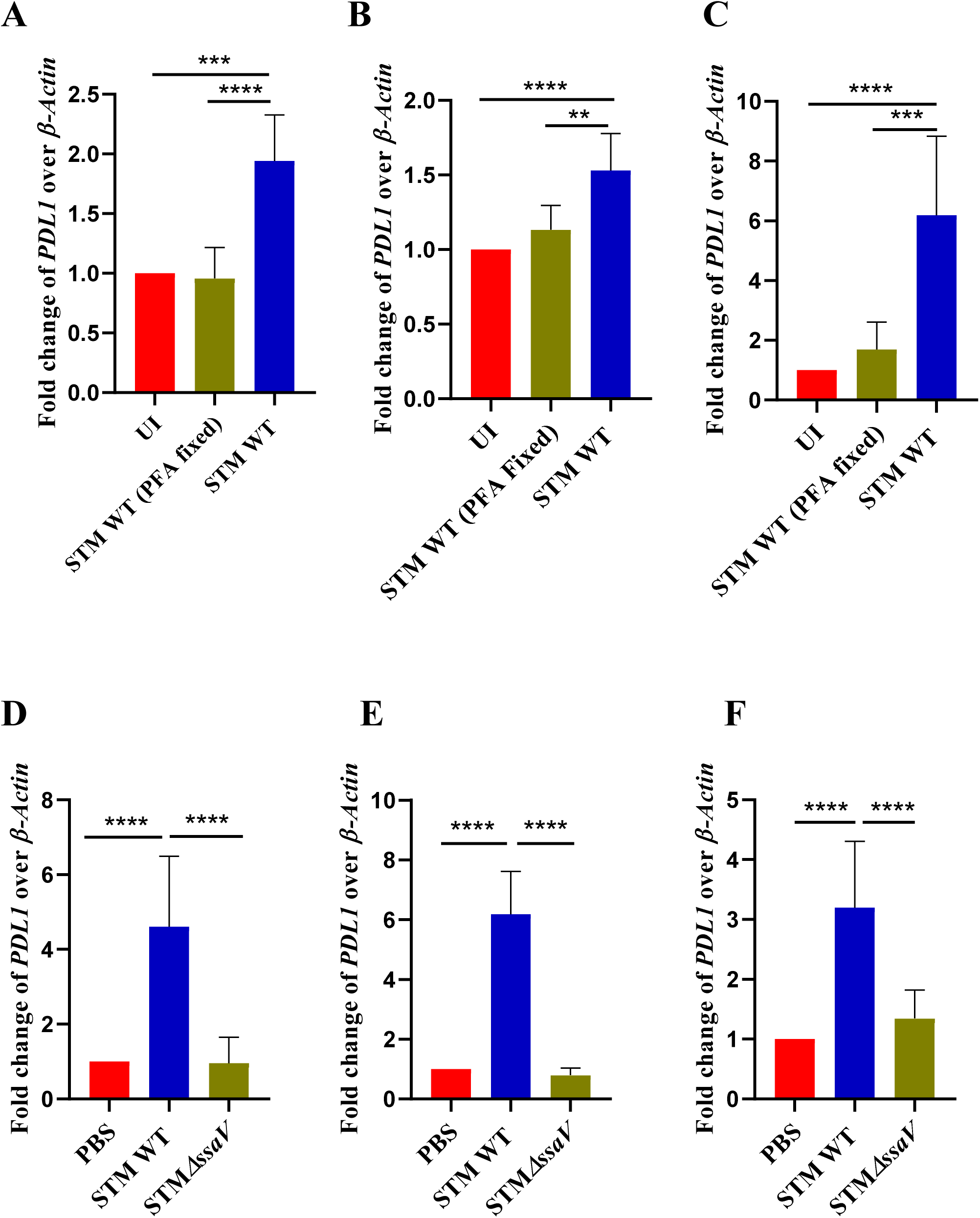
Salmonella upregulates the level of PD-L1 *in vitro* as well as *in vivo* using *Salmonella* pathogenicity island 2 (SPI-2) (A) Expression of PD-L1 was assessed using qRT-PCR in RAW 264.7 macrophages, 24 hours post-infection at MOI 10. Data is representative of N=3, n=3, presented as mean±SD. (B) Expression study of PD-L1 through qRT-PCR in bone marrow-derived macrophages (BMDM), 24 hours post-infection at MOI 10. Data is representative of N=2, n=3, presented as mean±SD. (C) Expression study of PD-L1 through qRT-PCR in Caco2 cell line, 24 hours post-infection at MOI 50. Data is representative of N=3, n=3, presented as mean±SD. (D) Expression study of PD-L1 through qRT-PCR in murine Liver (E) Spleen, (F) and Intestine, 5^th^ day post oral gavage. Data is representative of N=2, n=5 mice per cohort, presented as mean±SD. Statistical analysis was performed using a one-way ANOVA to determine p-values. (****p<0.0001, ***p<0.001, **p<0.01, *p<0.05)

### *Salmonella* effector SseL is crucial for upregulating PD-L1 *in vitro* as well *in vivo*

Figure 1 indicates that effector proteins from SPI2 are crucial for PD-L1 upregulation. Sahler *et al.,* also suggested the potential involvement of *Salmonella* effector SseL in PD-L1 upregulation in intestinal epithelial cell culture, but the mechanism behind it remains elusive [21]. We further validated the importance of SseL (a known deubiquitinase that targets NF-kB) in upregulating PD-L1 levels. We initiated the experiment by generating the knockout strains of *sseL* in the STM WT 14028s background. We first assessed the growth rate of the mutant strain in nutrient-rich LB media and observed no significant growth defects as compared to STM WT (Fig-S1A). To test the hypothesis that SseL is critical for PD-L1 upregulation, we infected the Caco2 cells with STM WT, STM*ΔssaV*, and STM*ΔsseL* at the MOI of 50 and harvested the cells 24 hours post-infection. We further quantified the transcript level of PD-L1 through qRT-PCR and observed where STM WT could upregulate PD-L1 levels in the Caco2 cell line, whereas STM*ΔssaV* and STM*ΔsseL* could not do so (Fig-2A). A similar observation was found through an immunofluorescence assay where STM WT infected Caco2 cells showed higher mean fluorescence intensity (MFI) or CTCF of PD-L1 than uninfected control, and STM*ΔsseL* infected, suggesting that changes in PD-L1 levels are evident at transcript as well as protein level (Fig-2 B, C). Further, to understand the importance of *Salmonella* effector SseL *in vivo*, we orally gavaged C57BL/6 mice, and, 5 days post-infection, quantified the transcript level of PD-L1 in mice organs like the liver and spleen. Quantification of data suggests where STM WT can upregulate the transcript level of PD-L1 in the mice organs like the liver and spleen, the STM*ΔsseL* infected cohort shows a reduced level of PD-L1 compared to STM WT (Fig-2D, E). Additionally, Western blot analysis corroborated these findings, revealing significantly lower PD-L1 levels in the STM*ΔsseL*-infected cohort compared to the STM WT-infected group (Fig-S1B, S1C). Taken together, this data highlights the importance of SseL in altering the levels of PD-L1 at transcript and protein levels *in vitro* and *in vivo*.

**Fig 2:**
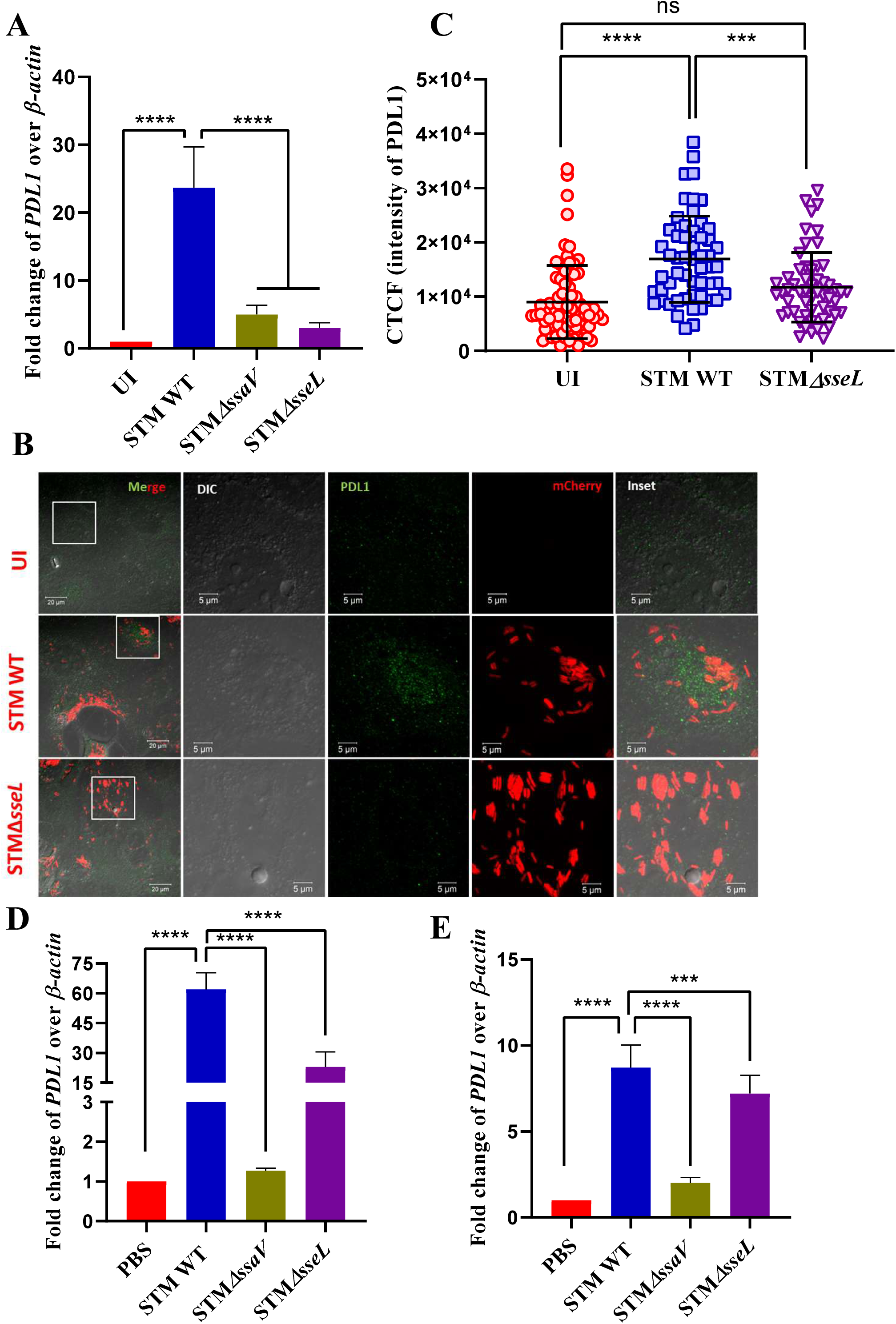
Salmonella effector sseL is crucial for upregulating PD-L1 *in vitro* as well as *in vivo*. (A) Expression study of PD-L1 through qRT-PCR in Caco2 cell line, 24h post-infection at MOI 50. Data is representative of N=3, n=3, presented as mean±SD. (B) Confocal microscopy images of uninfected and infected Caco2 cells, either with STM WT expressing mCherry or STM*ΔsseL* expressing mCherry, and immunostained against PD-L1 (green: Alexa fluor 488) (C) Quantification of cell total corrected fluorescence (CTCF) intensity of PD-L1 from the confocal images depicted in (B). Data is representative of n=80-100 cells from 3 independent experiments, presented as mean±SD. (D) Expression study of PD-L1 through qRT-PCR in murine liver, 5 days post-infection. Data is representative of N=3, n=5 mice per cohort, mean±SD. (E) Expression study of PD-L1 through qRT-PCR in murine spleen, 5 days post-infection. Data is representative of N=3, n=5 mice per cohort, mean±SD. Statistical analysis was performed using a one-way ANOVA to determine p-values. (****p<0.0001, ***p<0.001, **p<0.01, *p<0.05)

### STM*ΔsseL* shows a defect in colonization but induces higher inflammation

To investigate the pathogenicity and colonization of STM*ΔsseL in vivo*, we orally gavaged the mice with 10^7^ CFU per animal and assessed organ burden at 5th-day post-infection. Quantification of data suggests that compared to STM WT, there was a significant reduction in the colonization of STM*ΔsseL* in the liver and spleen, while STM*ΔssaV* was kept as a control (Fig-3A, B). To evaluate the importance of PD-L1 during the colonization of *Salmonella*, we performed an *in vivo* knockdown of PD-L1 using an adeno-associated virus (Fig-S2A, S2B) and subsequently infected the mice with 10^7^ CFU of STM WT. The data suggest that knocking down PD-L1 resulted in impaired colonization of STM WT in the liver and spleen (Fig-3C, D). Additionally, the cohort of mice treated with scrambled control lost more weight than the PD-L1 knockdown mice cohort following infection with STM WT (Fig-S2C), thus highlighting the importance of PD-L1 upregulation for *Salmonella* pathogenicity and colonization. Interestingly, despite having diminished colonization, the STM*ΔsseL* infected cohort exhibited earlier mortality (Fig-3E). Since the STM*ΔsseL* infected cohort shows a reduction in PD-L1 level compared to the STM WT-infected cohort, we hypothesized that the STM*ΔsseL*-infected cohort experienced heightened inflammation, contributing to earlier mortality despite lower organ burden. To test this hypothesis, we quantified the transcript level of pro-inflammatory genes such as IL-6 and TNF-α in murine liver. Quantification of data suggests that the STM*ΔsseL* infected cohort shows elevated transcript levels of inflammatory genes such as IL-6 and TNF-α (Fig-3F, G). In contrast, the transcript level of the anti-inflammatory gene IL-10, although not significant, shows a higher trend in the STM WT-infected cohort (Fig-3H). Further, we quantified the pro-inflammatory cytokine level of TNF-α and IL-6 in mice serum through ELISA and observed higher levels of these cytokines in the STM*ΔsseL* infected cohort (Fig-3I, J). To further elucidate the pathological differences, we performed hematoxylin and eosin (H&E) staining and quantified the number of infiltrated neutrophils in the liver. Upon quantification, we observed that the number of infiltrated neutrophils was not different in the STM WT and STM*ΔsseL* infected cohort (Fig-3K, L). However, we observed pronounced pathological changes in the liver sections infected with STM*ΔsseL,* characterized by several areas with neutrophil aggregation and large areas of necrosis compared to STM WT infected (Fig-3M). A well-defined and standard histopathological scoring method was employed with little modification (explained in materials and methods) to grade the inflammation/pathological score. Similarly, histopathology sections of intestines were analyzed, and upon quantification, we observed a high pathological score in the STM*ΔsseL* infected cohort (Fig-3N, O). Together, this data suggests that where STM*ΔsseL* shows defects in colonization in mice organs, but infected mice cohort demonstrated high-grade inflammation and succumbed to mortality earlier.

**Fig 3:**
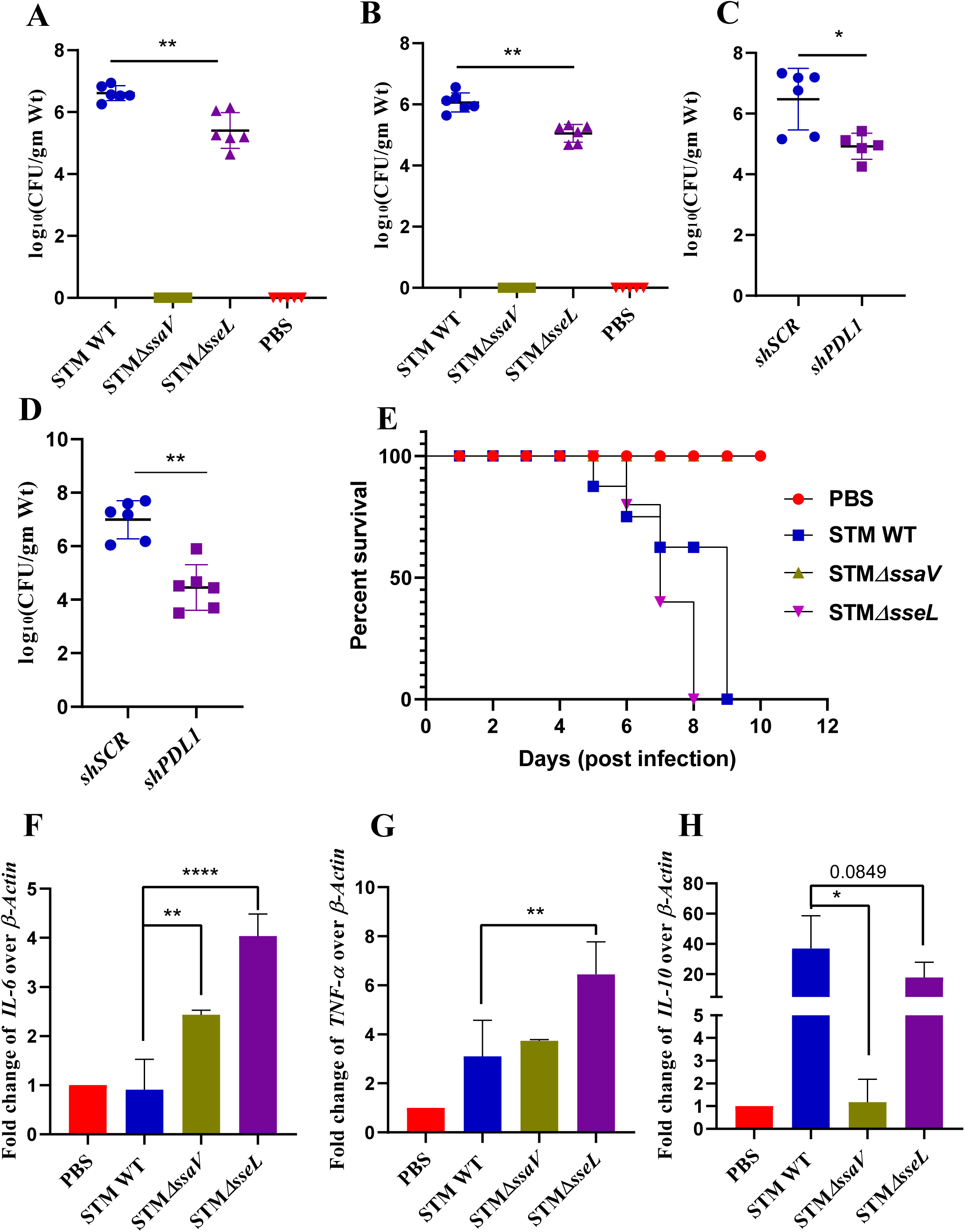
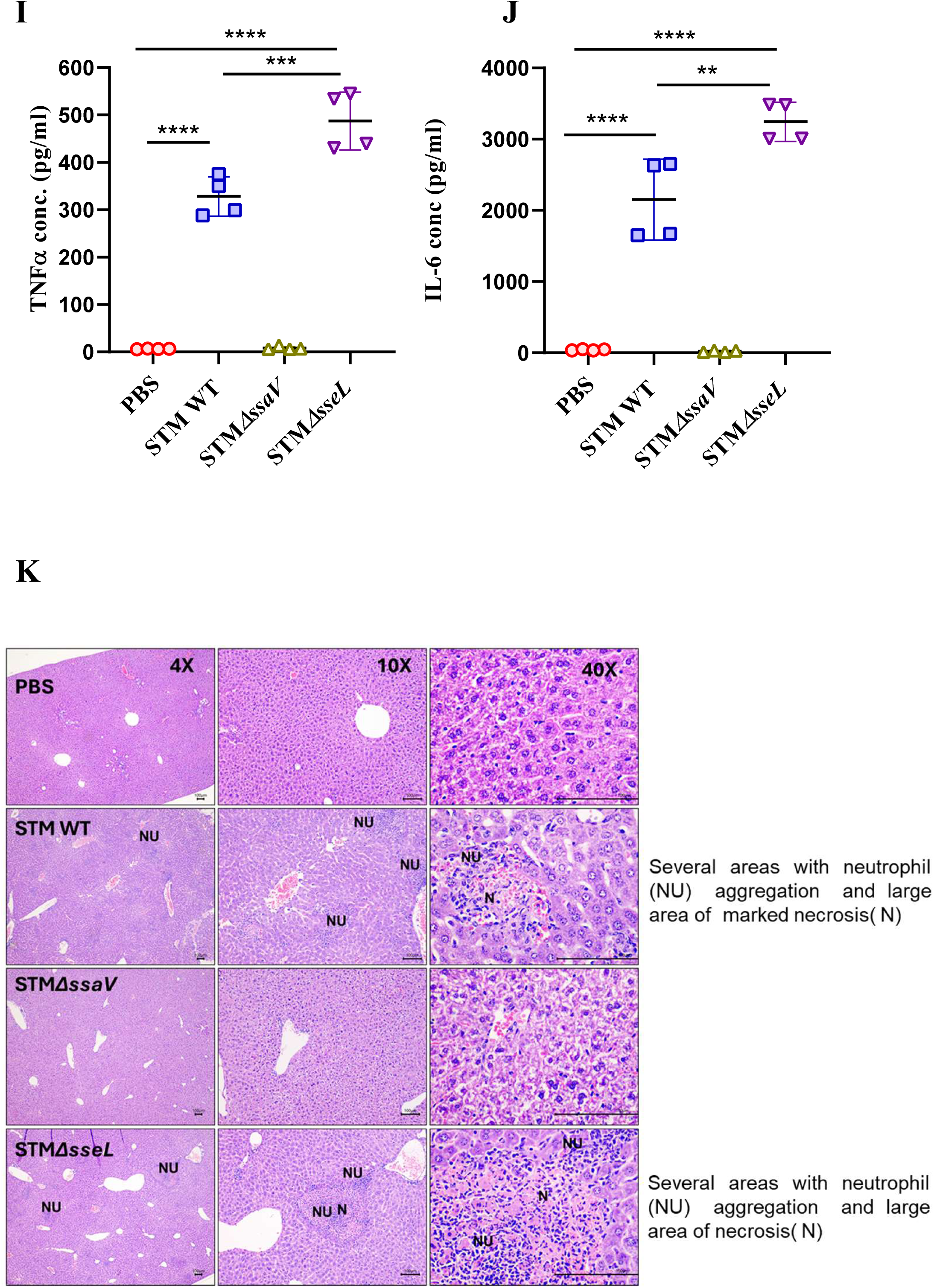
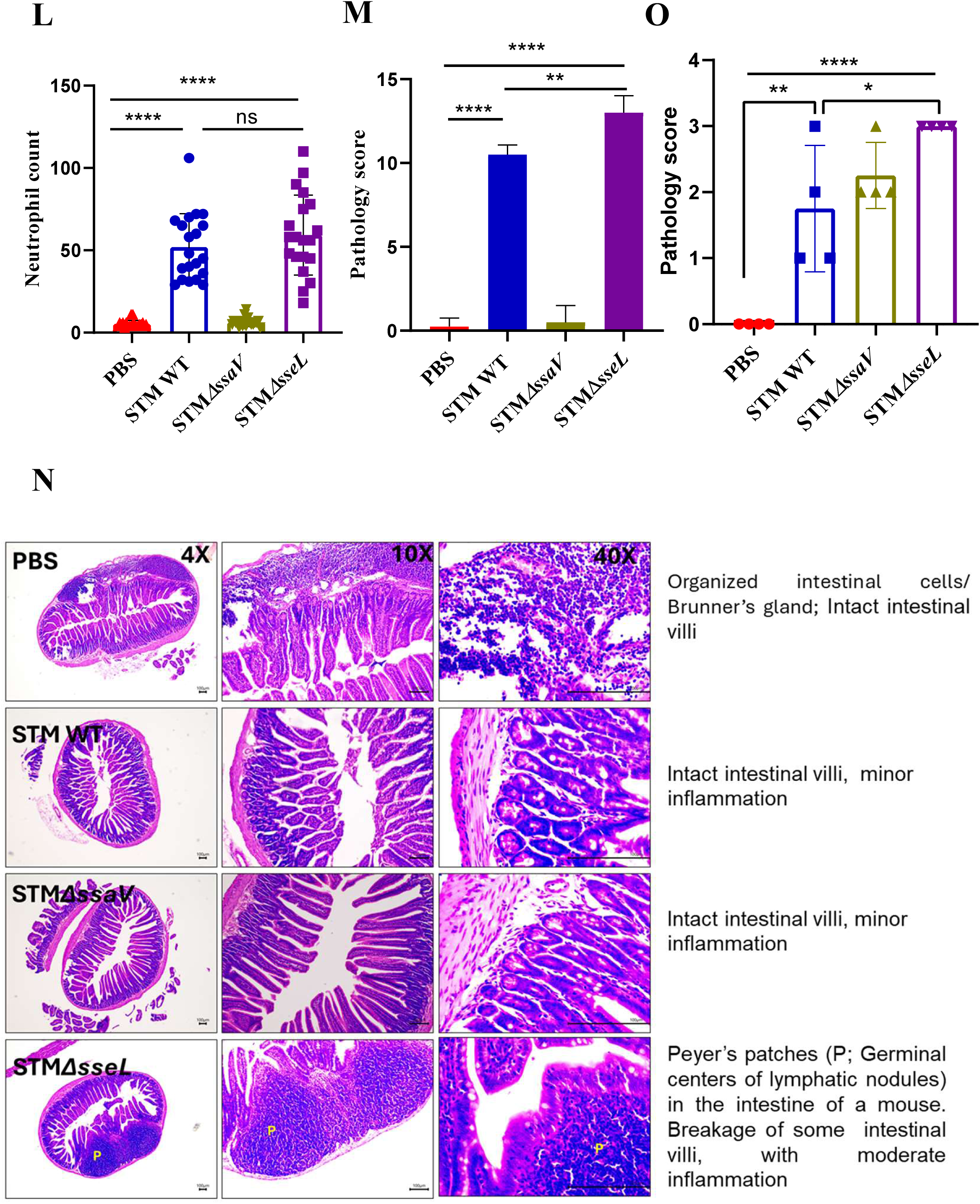
STMΔsseL shows a defect in colonization but induces higher inflammation. (A) *In-vivo* organ burden of STM WT, STM*ΔssaV,* and STM*ΔsseL* in the liver of C57BL/6 mice at day 5 post-infection. Data is representative of N=3, n=5 mice per cohort. Statistical analysis was conducted using the Mann-Whitney test to determine p-values (****p<0.0001, ***p<0.001, **p<0.01, *p<0.05). (B) *In-vivo* organ burden of STM WT, STM*ΔssaV,* and STM*ΔsseL* in the spleen from C57BL/6 mice at day 5 post-infection. Data is representative of N=3, n=5 mice per cohort. Statistical analysis was performed using the Mann-Whitney test for p-values (****p<0.0001, ***p<0.001, **p<0.01, *p<0.05). (C) *In-vivo* organ burden of STM WT in the liver of C57BL/6 mice at day 5 post-infection upon *in vivo* adenovirus-mediated PD-L1 knockdown. Data is representative of n=3 mice per cohort. Mann-Whitney Test was performed to obtain the p values. (****p<0.0001, *** p < 0.001, ** p<0.01, * p<0.05). (D) *In-vivo* organ burden of STM WT in the spleen from C57BL/6 mice at day 5 post-infection upon *in vivo* adenovirus-mediated PD-L1 knockdown. Data is representative of n=3 mice per cohort. Mann-Whitney Test was performed to obtain the p values. (****p<0.0001, *** p < 0.001, ** p<0.01, * p<0.05). (E) Percent survival of STM WT, STM*ΔssaV,* and STM*ΔsseL* infected C57BL/6 mice upon infection with 10^8^ CFU per animal. Data is representative of N=2, n=5 mice per cohort. (F) Expression of *IL6* was analyzed via qPCR in murine liver, 5 days post-infection. Data is representative of N=3, n=5 mice per cohort, presented as mean±SD. A one-way ANOVA test was performed to obtain the p values. (****p<0.0001, ***p < 0.001, ** p<0.01, *p<0.05) (G) Expression of *TNF-α* was assessed through qPCR in murine liver, 5 days post-infection. Data is representative of N=3, n=5 mice per cohort, presented as mean±SD. A one-way ANOVA test was performed to obtain the p values. (****p<0.0001, ***p < 0.001, ** p<0.01, *p<0.05) (H) Expression of *IL-10* was evaluated through qPCR in murine liver, 5 days post-infection. Data is representative of N=3, n=5 mice per cohort, presented as mean±SD. A one-way ANOVA test was performed to obtain the p values. (****p<0.0001, ***p < 0.001, ** p<0.01, *p<0.05) (I) Serum TNF-α levels of STM WT, STM*ΔssaV,* and STM*ΔsseL* infected C57BL/6 WT mice (males) mice at 5th day post-infection. Data is representative of N=2, n=2. A one-way ANOVA test was performed to obtain the p values. (****p<0.0001, ***p < 0.001, ** p<0.01, *p<0.05) (J) Serum IL-6 levels of STM WT, STM*ΔssaV,* and STM*ΔsseL* infected C57BL/6 WT mice (males) mice at 5th day post-infection. Data is representative of N=2, n=2. A one-way ANOVA test was performed to obtain the p values. (****p<0.0001, ***p < 0.001, ** p<0.01, *p<0.05). (K) Representative image of haematoxylin-eosin-stained liver sections to assess the liver tissue architecture upon *Salmonella* infection at 5^th^ days post-infection in various mice cohorts. (PBS treated, STM WT, STM*ΔssaV,* and STM*ΔsseL* infected). N=2, n=4, Scale bar-100µm.The scoring of necroinflammation is graded as 0-4; for each of the pathological lesions (Severe -4; Moderate -3; mild -2; Minor/minimum -1; No pathology -0) considering the following pathological lesions periportal hepatitis; focal inflammation, focal necrosis, and portal inflammation. (L) Quantification of neutrophil count in liver sections. (M) Graph representing the histopathological scoring of the liver sections depicted in (K). A one-way ANOVA test was performed to obtain the p values. (****p<0.0001, *** p < 0.001, ** p<0.01, * p<0.05). (N) Representative image of haematoxylin-eosin-stained intestine sections to assess the tissue architecture upon *Salmonella* infection at 5^th^ days post-infection in different mice cohorts. (PBS treated, STM WT, STM*ΔssaV,* and STM*ΔsseL* infected). Scale bar-100µm. (O) Graph representing the histopathological scoring of the intestine sections depicted in (N). Scoring system: The scoring of the pathological changes is graded as 0-4; for each of the pathological lesions (Severe/ marked -4; Moderate -3; mild -2; Minor/minimum -1; No pathology -0) considering the following pathological lesions, inflammation, tissue architecture. A one-way ANOVA test was performed to obtain the p values. (****p<0.0001, *** p < 0.001, ** p<0.01, * p<0.05).

### Deubiquitinase activity of SseL leads to β-catenin stabilization which directly regulates PD-L1 level

In unstimulated cells, transcription factor NF-kB is sequestered by IkB family proteins in the cytosol, preventing the translocation of NF-kB inside the nucleus. Activation of the NF-kB signaling pathway necessitates the degradation of IkB, allowing free NF-kB to translocate into the nucleus and regulate the expression of several pro-inflammatory genes. Previously, during *Salmonella* infection, SseL was reported to inhibit IkB degradation and NF-kB activation through its deubiquitinase activity, where in the absence of SseL, higher activation of NF-kB takes place [37]. This observation aligns with our findings as well where we have observed higher inflammation in STM*ΔsseL* infected mice cohort. Several studies have established that β-catenin signaling acts as a negative regulator of NF-kB mediating proinflammatory pathway [38].

Interestingly, both β-catenin and IkB are regulated and targeted for degradation by a similar mechanism. Specifically, both proteins are phosphorylated at N-terminal residues, followed by ubiquitination by the same ubiquitin ligase complex, E3-SCF^β-TrCP^, and further degraded through proteasome machinery [39]. Upon activation of β-catenin signaling, stabilized β-catenin translocates to the nucleus and, in conjunction with T-cell factor/ Lymphoid enhancer-binding factor (TCF/LEF), regulates the transcription of several downstream genes. Additionally, β-catenin signaling is known to regulate the level of PD-L1 in several cancers, glioblastoma, and viral infections [24], [25], [26], [34]. Based on existing literature, we hypothesize that the deubiquitinase activity of SseL can stabilize β-catenin during *Salmonella* infection, protecting it from degradation, and subsequently regulating PD-L1 level. To test the hypothesis if β-catenin mediated signaling is involved upon *Salmonella* infection, we infected the Caco2 cells with STM WT and STM*ΔsseL* while keeping uninfected cells and STM*ΔssaV* infected as a control and quantified the expression of downstream target genes. Quantification of data suggests that upon STM WT infection, there is an increase at the transcript level of downstream target genes such as Axin2 and CyclinD, while no difference was observed upon STM*ΔsseL* infection compared to uninfected cells, suggesting in the absence of SseL, β-catenin signaling is not active (Fig-4A, B). We have also observed an increase in the N-terminal phosphorylated state of β-catenin (enhanced N-terminal phosphorylation leads to faster degradation) upon infection with STM*ΔsseL* (Fig-4C, D). Additionally, we observed increased mean fluorescence intensity (MFI) or CTCF of β-catenin inside the nucleus of STM WT-infected Caco2 cells as compared to STM*ΔsseL* infected and uninfected cells (Fig-4E, F). Till now our data suggests that *Salmonella* regulates β-catenin signaling and activates the expression of several target genes. In the case of glioblastoma, β-catenin directly regulates the level of PD-L1 by binding to its promoter region [34]. This finding prompted us to investigate whether β-catenin similarly regulates PD-L1 expression during *Salmonella* infection by binding to its promoter. To understand the same, we performed a ChIP assay and observed an increase in enrichment of β-catenin recruitment on the promoter of PD-L1 (gene: CD274) upon STM WT infection. In contrast, β-catenin enrichment was significantly lower in STM*ΔsseL*-infected cells, although it was recovered to WT levels upon complementation of the *sseL* gene in the STM*ΔsseL* background (Fig-4G). To further elucidate the role of β-catenin in this upregulation, we further treated the cells with β-catenin inhibitor (FH535) and quantified the transcript level of PD-L1. Data indicate that while STM WT can upregulate PD-L1 levels in DMSO-treated conditions compared to STM*ΔsseL*, PD-L1 levels do not increase upon STM WT infection when treated with the inhibitor FH535 (Fig-4H). Additionally, a TOP flash luciferase reporter assay was conducted to assess β-catenin activation in the nucleus. Data suggests that compared to STM WT, there is significantly less luciferase activity upon infection with STM*ΔsseL* (Fig-S3A). Taken together, this data suggests that *Salmonella* effector SseL regulates the β-catenin and further stabilized β-catenin translocates into the nucleus and directly regulates the level of PD-L1.

**Fig 4:**
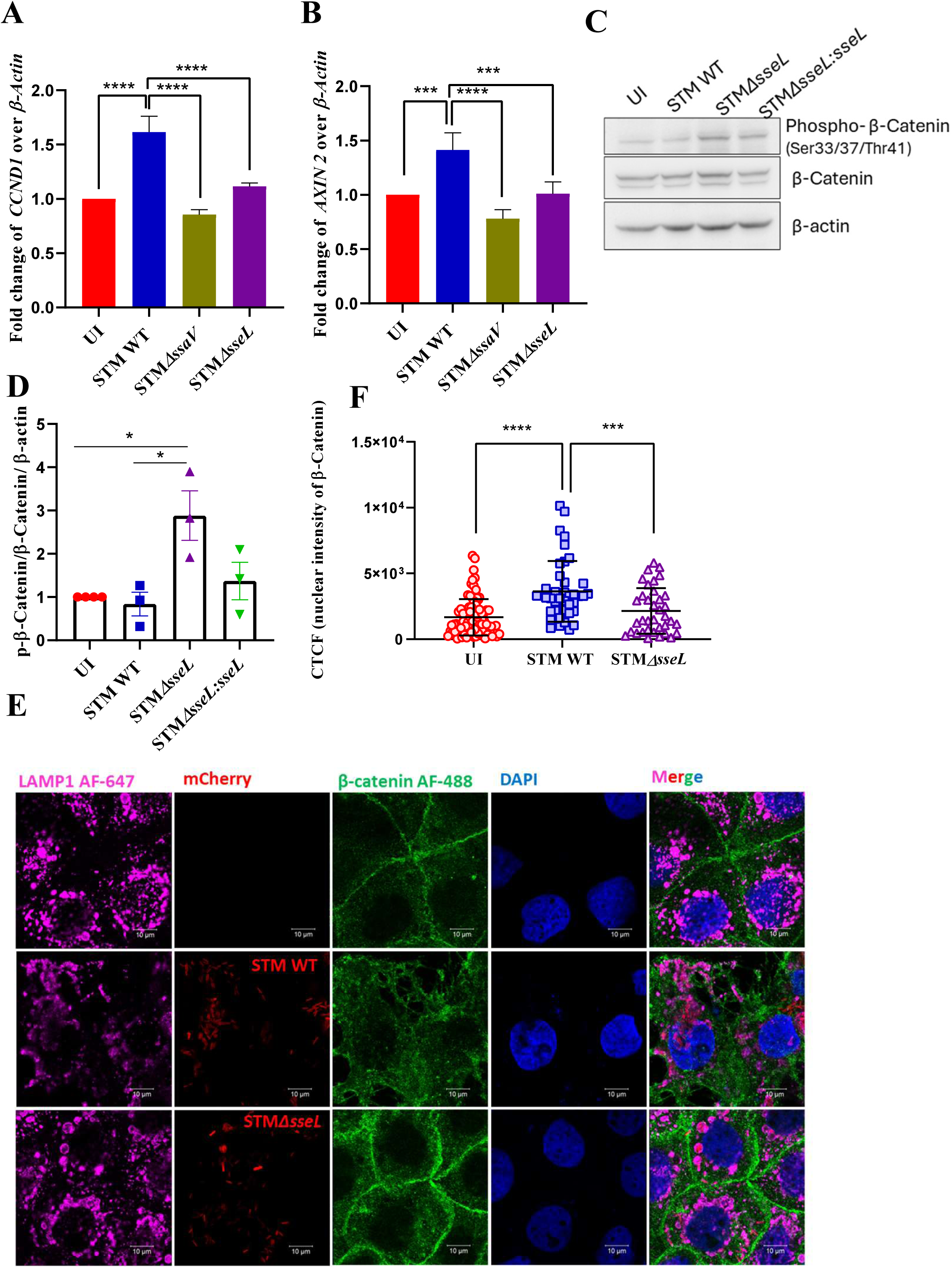
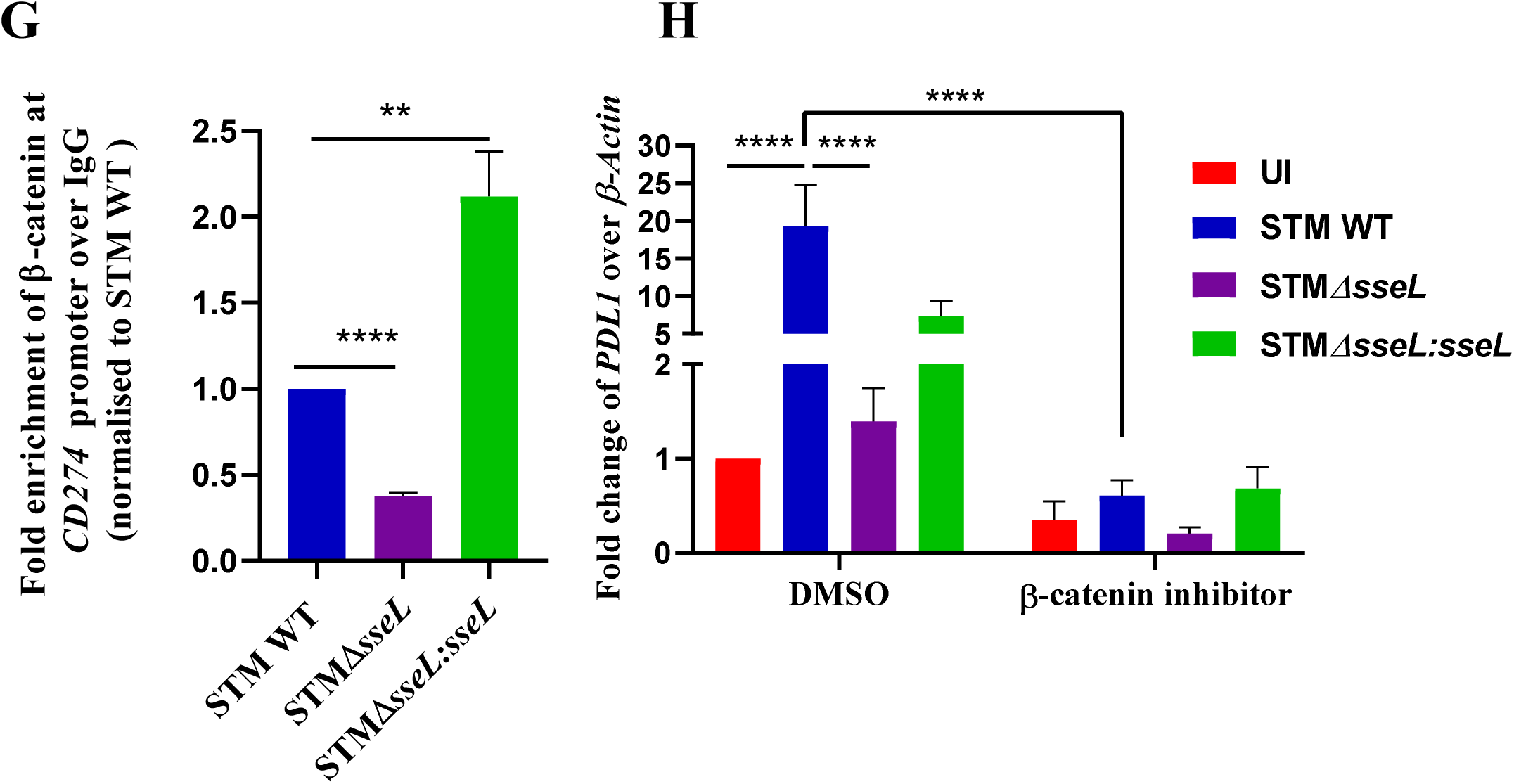
Deubiquitinase activity of SseL leads to β-catenin stabilization which directly regulates PD-L1 level. (A) Expression study of *CCND1* through qPCR in Caco2 cells, 24 hours post-infection. Data is representative of N=3, n=3, presented as mean±SD. A one-way ANOVA test was performed to obtain the p values. (****p<0.0001, ***p < 0.001, ** p<0.01, *p<0.05). (B) Expression study of *AXIN2* through qPCR in Caco2 cells, 24 hours post-infection. Data is representative of N=3, n=3, mean±SD. A one-way ANOVA test was performed to obtain the p values. (****p<0.0001, ***p < 0.001, ** p<0.01, *p<0.05). (C) Immunoblot of phospho-β-catenin and total β-catenin in uninfected, STM WT, STM*ΔsseL* and STM*ΔsseL:sseL* infected Caco2 cells, 24 hours post-infection. (D) Densitometric plot depicting the band intensities of phospho-β-catenin normalized to total β-catenin and loading control, β-actin from blot (C), data is representative of N=3, presented as mean±SEM. A one-way ANOVA test was performed to obtain the P-values. (****p<0.0001, ***p < 0.001, ** p<0.01, *p<0.05). (E) Confocal microscopy images of uninfected and infected Caco2 cells with either STM WT expressing mCherry or STM*ΔsseL* expressing mCherry, immunostained against LAMP1 (Alexa fluor-647) and β-catenin ( Alexa fluor-488) (F) Quantification of cell total corrected fluorescence (CTCF) intensity of β-catenin inside the nucleus from the confocal images depicted in (E). Data is representative of n=90-100 cells from 3 independent experiments, presented as mean±SD. A one-way ANOVA test was performed to obtain the p values. (****p<0.0001, ***p < 0.001, ** p<0.01, *p<0.05). (G) Chromatin immunoprecipitation (ChIP) assays using anti-IgG and anti-β-catenin antibodies in STM WT, STM*ΔsseL*, and STM*ΔsseL:sseL* infected Caco2 cells. Quantitative PCR analyses were conducted with primers specific to the promoter region of CD274. Statistical significance was assessed using a one-way ANOVA (****p<0.0001, ***p<0.001, **p<0.01, *p<0.05). (H) Expression study of PD-L1 through qPCR in Caco2 cells, 24 hours post-infection, with DMSO or inhibitor treatment (15µM). Data is representative of N=3, n=3, presented as mean±SD. A one-way ANOVA test was performed to obtain the p values. (****p<0.0001, ***p < 0.001, ** p<0.01, *p<0.05).

### An increase in the level of PD-L1 upon *Salmonella* infection leads to T cell inactivation

Since we have observed that using effector SseL, *Salmonella* upregulates the expression level of PD-L1, while an increase in PD-L1 is known to negatively regulate the T cell activity. To understand the functional importance of this increase upon infection, we performed a coculture experiment between infected Caco2 and Jurkat T cells. Briefly, Caco-2 cells were infected with STM WT, STM*ΔsseL*, or the complemented strain STM*ΔsseL:sseL* for 24 hours. Following infection, activated Jurkat T cells (stimulated with ionomycin and PMA) were added to the infected Caco-2 cells and co-cultured for an additional 12 hours. Subsequent analysis of Jurkat T cells revealed a significant increase in pro-inflammatory cytokines, specifically *IFN-γ* and *IL-1β*, when co-cultured with STM*ΔsseL*-infected Caco-2 cells compared to those infected with STM WT or STM*ΔsseL:sseL*. These results suggest that the elevated PD-L1 levels during STM WT infection suppress T cell cytokine production and activity. Conversely, the lack of PD-L1 upregulation in STM*ΔsseL* infection correlates with enhanced T cell activation and increased transcript levels of pro-inflammatory cytokines (Fig-5A, B). Overall, this data highlights the relevance and importance of PD-L1 upregulation upon *Salmonella* infection and this increase in PD-L1 level negatively regulates T-cell activity.

**Fig 5:**
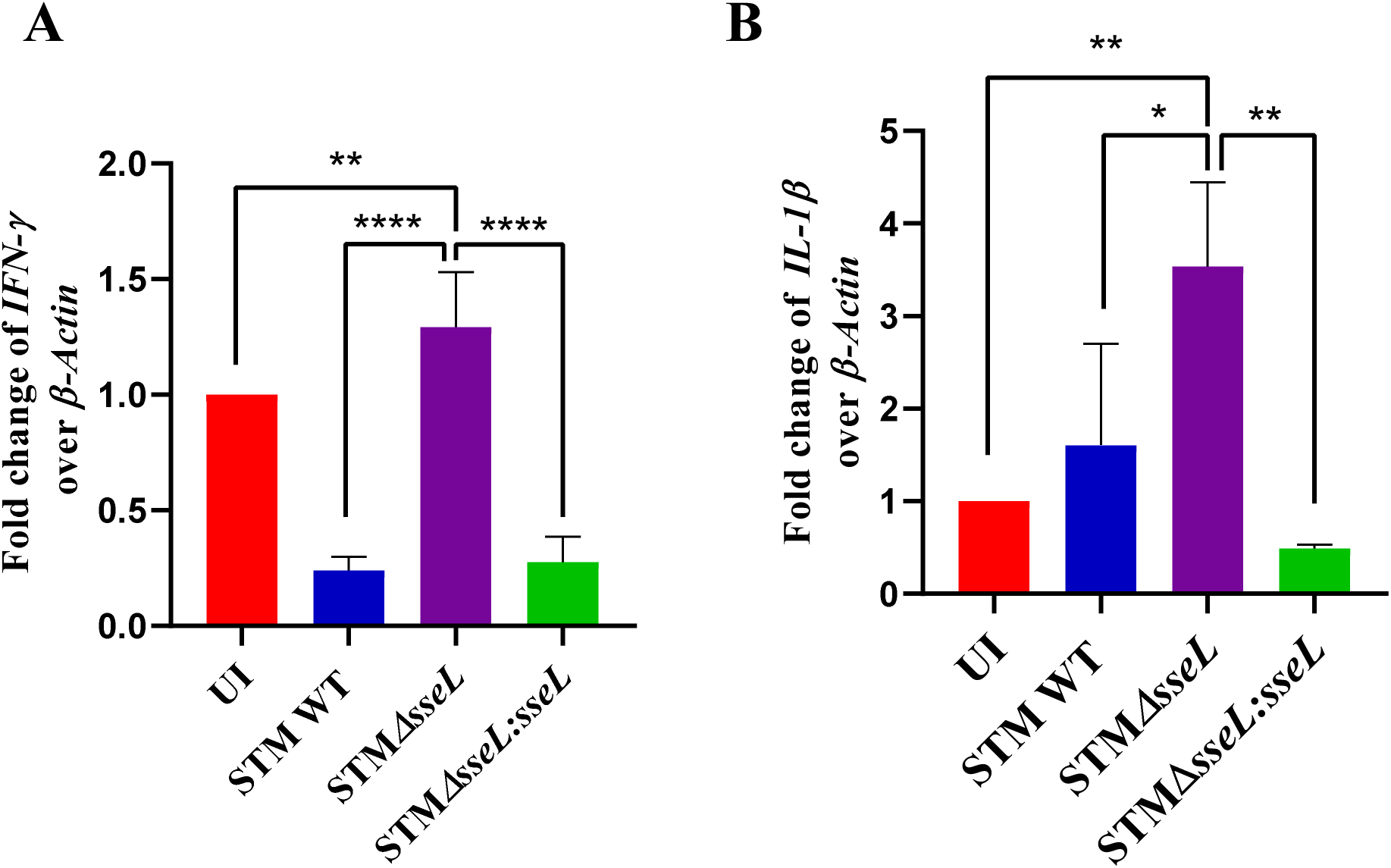
An increase in PD-L1 levels upon *Salmonella* infection leads to T cell inactivation. (A) Expression study of *IFNγ* through qPCR in Jurkat cells, 12 hours post-co-culture with infected Caco2 cells; Data is representative of N=3, n=3, presented as mean±SD. A one-way ANOVA test was performed to obtain the p values. (****p<0.0001, ***p < 0.001, ** p<0.01, *p<0.05). (B) Expression study of *IL-1β* through qPCR in Jurkat cells, 12 hours post-co-culture with infected Caco2 cells; Data is representative of N=3, n=3, presented as mean±SD. A one-way ANOVA test was performed to obtain the p values. (****p<0.0001, ***p < 0.001, ** p<0.01, *p<0.05).

## Discussion

Numerous pathogens use an arsenal of effector proteins to inhibit or evade the host immune response to establish a successful infection. *Salmonella,* being a successful pathogen, employs various strategies to evade both innate and adaptive immune responses. Among these strategies are the inhibition of the fusion of SCV with lysosomes, reducing the number of acidic lysosomes within the host cell, inhibiting xenophagy, polarizing M1 (pro-inflammatory) macrophages to M2 (anti-inflammatory) type, downregulation of the surface MHC-Ⅱ expression, and upregulating the level of immune checkpoint inhibitor such as PD-L1 [15], [21], [40], [41], [42]. In this study, we investigated to understand the molecular mechanism involved in PD-L1 upregulation. Our findings indicate that the upregulation of PD-L1 is facilitated by bacterial effector SseL, highlighting this as an active and stealthy strategy employed by *Salmonella* to subvert host immune defenses.

As reported by Sahler *et al*. regarding the significance of SseL in PD-L1 upregulation in intestinal epithelial cell lines, our results indicate that the *Salmonella* effector SseL is essential for this upregulation, both in cell culture and in animal models. Bacterial pathogens such as *Helicobacter pylori* and *Streptococcus* also upregulate PD-L1 through the action of effector protein CagA and surface protein PspA, respectively [24], [33]. Additionally, several viral pathogens, including Hepatitis B virus (HBV) also lead to PD-L1 upregulation [27]. These effectors primarily target cellular pathways, directly or indirectly regulating the expression of immune-inhibitory molecules. This evidence suggests that, through evolution, pathogens have acquired a diverse array of virulence factors to effectively modulate the host’s adaptive immune response.

*Salmonella* also downregulates the level of MHC-Ⅱ complex using effector SteD [14]. In this present study, we report an additional mechanism for PD-L1 upregulation by *Salmonella* through the effector SseL, which collectively serves to negatively regulate T cell proliferation. In our *in vivo* infection model, the cohort infected with STM*ΔsseL* exhibited significantly reduced levels of PD-L1 alongside an increase in pro-inflammatory cytokines. This heightened inflammatory response led to earlier mortality in this cohort compared to those infected with STM WT, highlighting the importance of PD-L1 upon infection. However, during *in vivo* pathogenesis, multiple factors are involved to modulate the immune system. In the STM*ΔsseL*-infected cohort, the observed high levels of inflammation may also result from additional signalling pathways alongside the PD-L1 axis [43]. Notably, STM*ΔsseL* demonstrated diminished colonization in mouse organs compared to STM WT. Furthermore, Geng et al. (2019) highlight the significance of SseL in regulating inflammatory cytokines in a distinct chicken model infected with *Salmonella pullorum*, reinforcing the broader implications of SseL’s role in immune modulation[44].

In this study, we observed the role of SseL in activating WNT/β-catenin signaling, where we observed more N-terminal phosphorylation of β-catenin in absence of SseL. In case of STM WT infection, stabilized β-catenin translocates to nucleus and regulates the level of PD-L1. A similar strategy is employed by the Hepatitis B virus (HBV), which activates PTEN/β-catenin/c-Myc signalling through its proteins HBx and HBp, leading to the upregulation of PD-L1 [27]. The changes in PD-L1 level directly regulate the T cell response, indicating that pathogens target the WNT/β-catenin pathway to modulate host immune responses for their proliferation and survival. Additionally, AvrA, another *Salmonella* effector protein with deubiquitinase activity, plays a crucial role in pathogenesis by inhibiting NF-κB activation and activating the β-catenin pathway [45], [46], [47], [48]. While SseL was known to deubiquitinate either IkB or ribosomal protein S3 (RPS3) [43], ultimately affecting NF-kB-mediated gene transcription, we report here for the first time that SseL also modulates the β-catenin pathway. This suggests that SseL may impact other cellular pathways, further influencing host immune responses. Moreover, investigating the pathogenesis of *Salmonella* strains with simultaneous knockout of both *sseL* and *avrA* would provide valuable insights into their cooperative roles. Given that these deubiquitinases are specific to *Salmonella*, they present promising targets for the development of novel therapeutic strategies.

## Acknowledgements

We acknowledge the Departmental and Divisional Confocal Microscopy Facility and Central Animal Facility at IISc for providing us with an opportunity for experimentation. Mr. Sumitlal K have helped in image acquisition in Confocal Microscopy. We acknowledge Prof. Kumaravel Somasundaram, IISc for providing TOPFlash plasmid.

## Author Contributions

UC and DC have conceived the study and designed the experiments. UC and MKS have performed all the experiments and participated in acquiring the data. UC and MKS have analyzed the data. UC has constructed the figures and wrote the original draft of the manuscript. UC, MKS, and DC have participated in the proofreading and editing of the manuscript. RSR has performed the histopathological scoring of the liver sections and the retro-orbital injection of the adenoviral constructs in mice. ADC and SKG have prepared and provided AAV2, for *in vivo* mice knockdown experiments. DC has contributed to the funding acquisition, project administration, and overall supervision of the work. All authors approved the final version of the manuscript.

## Legends to supplementary figure

**Supplementary Fig-1: *Salmonella* effector SseL is crucial for upregulating PD-L1 *in vitro* as well as *in vivo***

(A) Growth curve of STM WT and STM*ΔsseL* strains in nutrient-rich LB media over a 12-hour period by measuring optical density (OD) at 600nm. (B) Immunoblot of PD-L1 from the murine liver lysate of PBS-treated mice, and infected with STM WT, STM*ΔssaV*, and STM*ΔsseL*, collected on 5^th^-day post oral gavage. Data is representative of N=2, n=2. (C) Densitometric plot depicting band intensities of PD-L1 and normalized to loading control, β-actin in blot (B), presented as mean±SD. A one-way ANOVA test was performed to obtain the p values. (****p<0.0001, ***p < 0.001, ** p<0.01, *p<0.05).

**Supplementary fig-2: STM*ΔsseL* shows a defect in colonization but induces higher inflammation**

(A) Immunoblotting of PD-L1 in harvested liver samples upon AAV-mediated scrambled (*shSCR*) treated or *in vivo* PDL1knockdown (*shPDL1*) in mice, Data is representative of n=3. (B) Densitometric plot depicting the band intensities of PD-L1 normalized to loading control, β-actin in blot (A), data is representative of n=3, presented as mean±SD. (C) Weight reduction in mice: uninfected or infected with STM WT upon *in vivo* knockdown of PD-L1 or scrambled control.

**Supplementary fig-3 Deubiquitinase activity of SseL leads to β-catenin stabilization which directly regulates PD-L1 level**

(A) β-catenin luciferase activity in Caco2 cells upon infection with either STM WT or STM*ΔsseL*, 24 hours post-infection. Data is representative of N=2, n=3, presented as mean±SD. A one-way ANOVA test was performed to obtain the p values. (****p<0.0001, ***p < 0.001, ** p<0.01, *p<0.05).

