## Supplementary figures for "*Salmonella* effector SseL induces PD-L1 up-regulation and T cell inactivation via β-catenin signalling axis"

**Supplementary Fig-1:** ***Salmonella*** **effector sseL is crucial for upregulating PDL1** ***in vitro*** **as well as** ***in vivo***

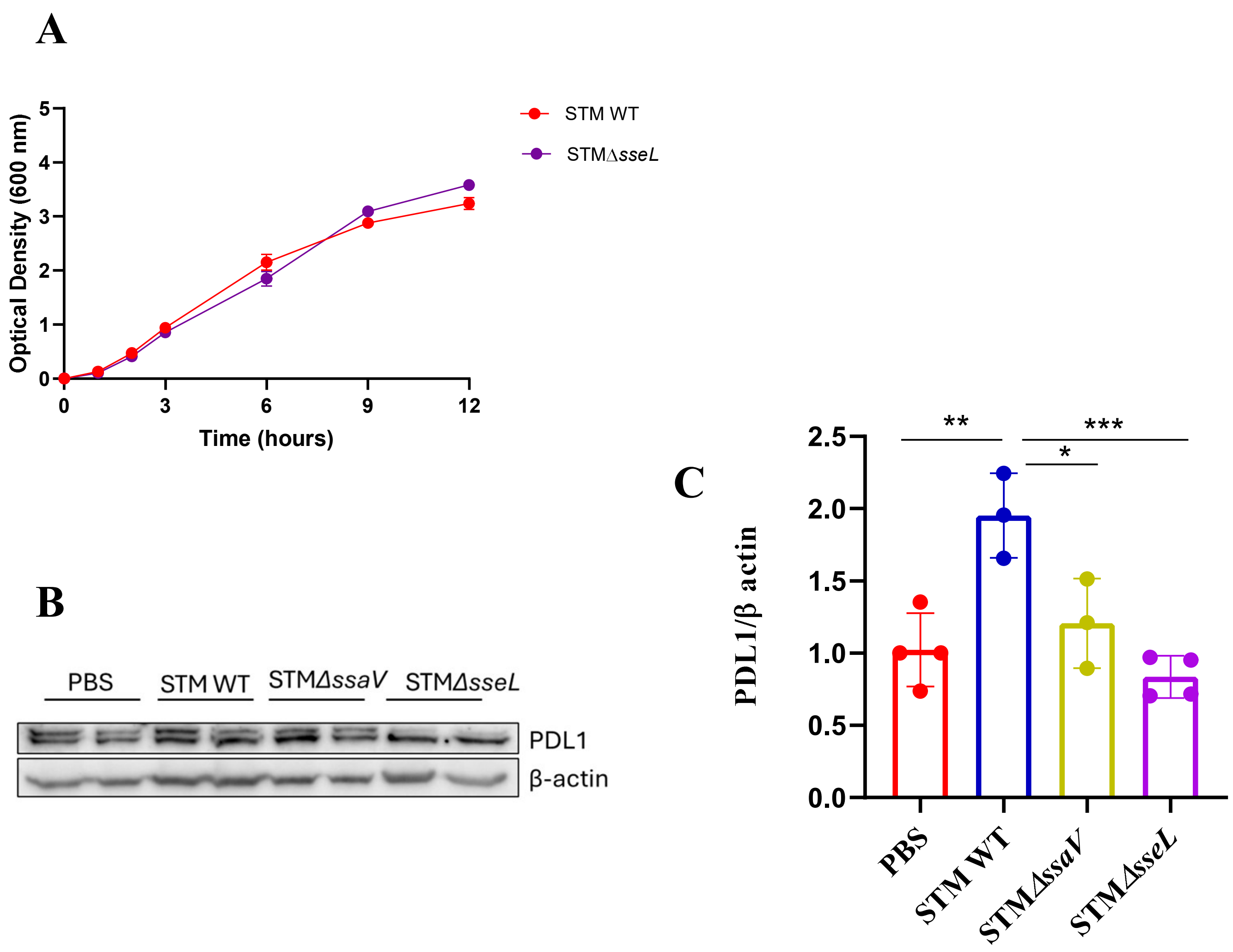

**Supplementary fig-2: STM*ΔsseL* shows a defect in colonization but induces higher inflammation**

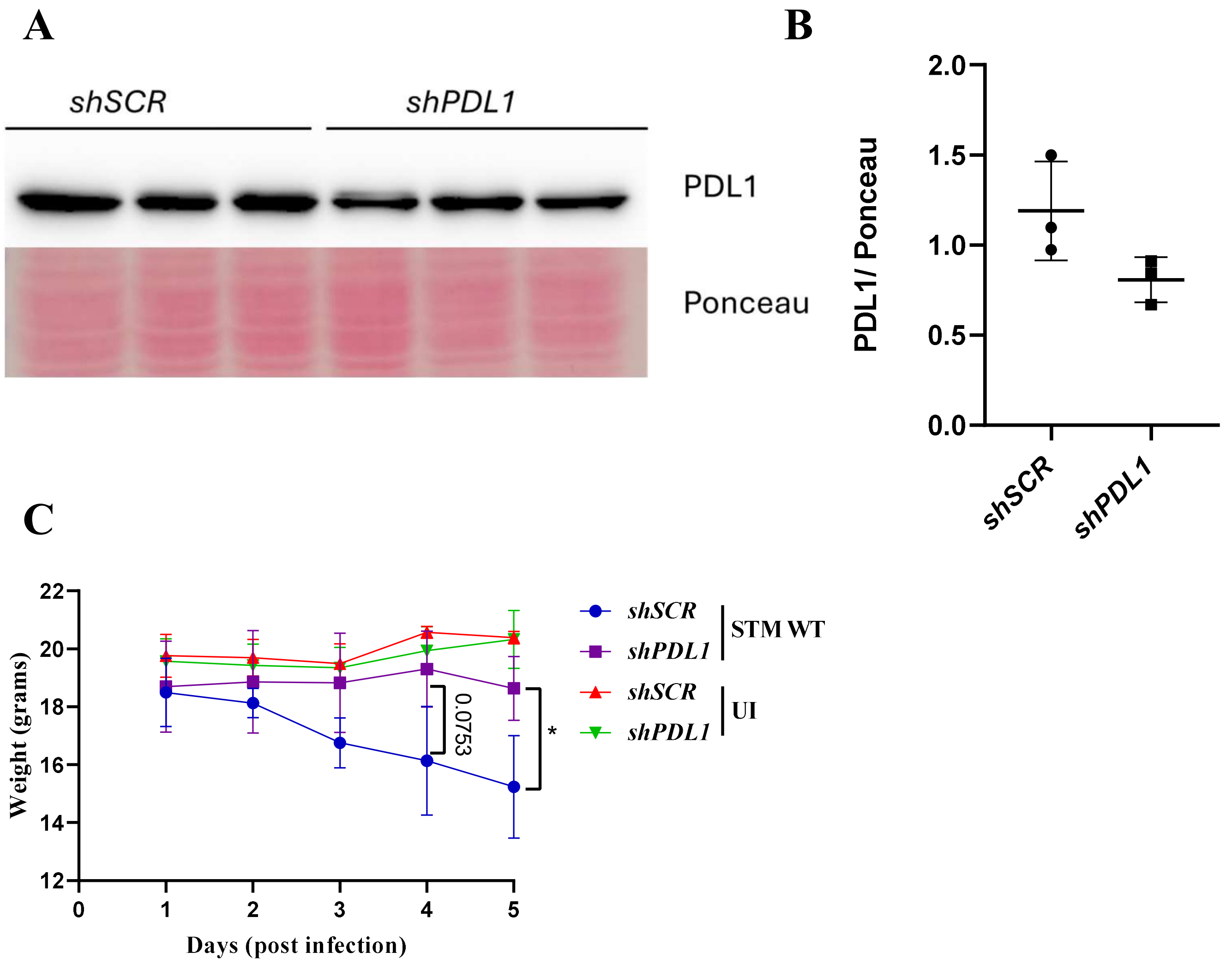

**Supplementary fig-3 Deubiquitinase activity of SseL leads to β-catenin stabilization which directly regulates PDL1 level**

**A**

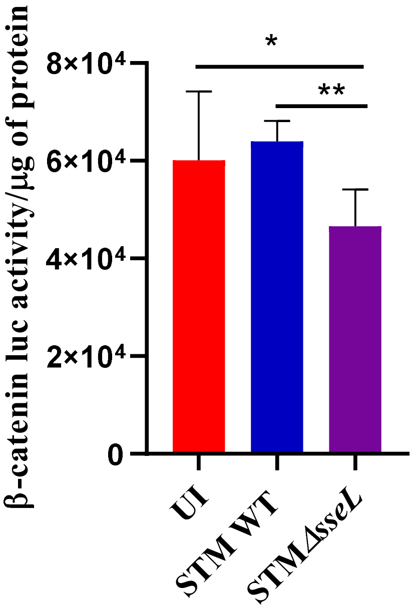
